# Effects of feeding treatment on growth rate and performance of primiparous Holstein dairy heifers

**DOI:** 10.1101/760082

**Authors:** Yannick Le Cozler, Julien Jurquet, Nicolas. Bedere

## Abstract

The objective of this study was to investigate effects of feeding-rearing programs that aim for first calving at 20-27 months (mo) of age on growth, reproduction and production performance of Holstein cows at nulliparous and primiparous stages. We hypothesised that, in a seasonal autumn-calving strategy, heifers born late in the season could catch up to the growth of heifers born earlier and be inseminated during the same period, at a body weight (BW) of at least 370 kg. This approach would result in first calving age at 21-22 mo of age without impairing their later performance. To test this hypothesis, we studied 217 heifers over 3 years. They were split into three treatment groups: control feeding (SD), an intensive-plane diet (ID1) from birth to 6 mo of age or an intensive-plane diet from birth to one year of age. Heifers in groups SD and ID1 were born from September until the end of November, while those in ID2 were born later. The present study showed that late-born heifers (ID2) could catch up with the growth of the others due to the feeding treatment, although they were still 42 kg lighter than the SD and ID1 heifers at first calving. No difference in reproductive performance was observed among groups. Once primiparous, the cows reared with the ID2 treatment tended to produce less milk than SD and ID1 cows (*ca*. 400 kg less on a 305 d basis throughout lactation), and no differences in milk composition, feed intake, body condition score or BW were observed among groups. Age at first service (AFS) was classified *a posteriori* into three classes: 12.5 (AFS_12.5_), 14.0 (AFS_14.0_) and 15.5 mo (AFS_15.5_) of age. Heifers in AFS_12.5_ grew faster than those in AFS_14.0_ and AFS_15.5_. Once primiparous, the AFS_12.5_ cows tended to produce less milk at peak than AFS_14.0_ and AFS_15.5_ cows (*ca*. 1.5 kg/d less) although no difference in total milk yield during lactation was observed. No differences in milk composition, feed intake, body condition score or BW were observed among groups. These results support the conclusion that the feeding treatment can enable late-born heifers to catch up to the growth of heifers born earlier in the season. This strategy results in an earlier first calving that does not impair their reproductive performance but does decrease milk yield slightly during first lactation. Future studies should investigate long-term effects of this strategy.

## Introduction

In seasonal calving systems, heifers usually first calve at a young age (*ca*. 24 months: mo). The first insemination (*i.e*. service) may be delayed, however, for heifers born at the end of the calving period if an adequate body weight (BW) is not reached (*i.e*. 360-380 kg for Holstein heifers in French dairy herds; Le Cozler *et al*., 2008). Increasing nutrient uptake and thus the growth rate of these late-born heifers is one solution to lower this risk. High growth rate during rearing is associated with decreased age at puberty; consequently, first calving may occur as early as 20-21 mo of age. Tozer (2000) concluded that a higher plane of nutrition incurred higher daily feed costs, but these costs were recouped when heifers calved at a younger age through savings on labour, housing and overall feed costs. Regardless of the rearing strategy (group-calving or not), animals need to reach an adequate body size and or body weight before calving to avoid compromising milk production during the first lactation (Bach and Ahedo, 2008). Indeed, an accelerated growth program for dairy heifers cannot focus only on early onset of puberty. Many authors have studied the influence of growth intensity on future performances (Le Cozler *et al*., 2008). Most studies indicated that a too-rapid growth rate had a negative influence, while some indicated that accelerated growth had little impact. According to Pirlo *et al*. (1997), reducing the age of first calving to 23 to 24 mo was the most profitable procedure, but no less than 22 mo (except in cases of low milk prices and high rearing costs). They concluded that the reluctance to decrease the age of first calving is generally attribute to the belief that early calving is detrimental to milk yield and longevity. We designed and conducted an experiment to determine the influence of feeding treatments on growth parameters, reproduction and the production performance of Holstein primiparous heifers that first calved from 20-27 mo of age in a seasonal calving system. We assumed that genetic improvements in dairy production over the past few decades had yielded animals that could calve earlier than 24 mo of age. We also assumed that results for animals reared in a seasonal calving strategy could be used and generalised for those in a non-grouped strategy. We examined the potential for late-born heifers to catch up to the rest of the heifers by the first artificial insemination (AI) at a minimum BW of 370-380 kg, resulting in a first calving at less than 22 mo of age.

## Materials and Methods

### General design

A total of 217 Holstein heifers, born during the calving season in 2009-10 (n = 65), 2010-11 (n = 73) and 2011-12 (n = 76; September to February), were reared and followed until oestrus synchronisation (12-15 mo of age) at the INRA experimental farm of Méjusseaume (Le Rheu, France). For details of the rearing procedures and strategies used in the present study, see Abeni *et al*. (2019). Calves born from 1 September to 30 November were alternately assigned to 1 of two nutritional treatments (according to birth order) and fed either a standard diet (SD) or an intensive-plane diet (ID1) from 0-6 mo of age. It was expected that heifers fed the SD and ID1 diets would reach 190-200 and 220-230 kg at 6 mo of age, respectively. Heifers born after 1 December (ID2) received the same intensive-plane diet as ID1 heifers from 0-6 mo of age to decrease potential interaction between age and treatment during this period. Thereafter, a supplemental diet was formulated for ID2 heifers to enable them to reach 380 kg at 12 mo of age. The main objective of the ID2 diet was to study the potential for late-born heifers to catch up to the rest of the heifers by the first AI at a minimum BW of 370-380 kg. It was expected that this strategy would correspond to a mean age of 15 mo for SD and ID1 heifers and 12 mo for ID2 heifers. In year one, heifers grazed from mid-May until the end of October. In year two, heifers grazed from the end of March until calving season (starting 1 September). At the end of the first grazing season, all heifers were group-housed until being turned out to pasture in the second season. Three weeks before the expected date of calving, heifers were placed in cow herds and individually fed a similar total mixed ration (TMR). During lactation, milk yield was recorded twice per day and animals were weighed one per day. The experiment ended 15 weeks after calving.

### Feeding management

Diets were formulated for each growth stage according to recommendations and procedures developed by Agabriel and Mechy (2007) to reach a targeted average daily gain (ADG) per period, as a function of the initial BW and feeding treatment used. In this approach, energy is expressed per UFL (forage unit for lactation, UFL/kg dry matter: DM), which is the energy required for lactation (g/kg)/1760. For protein, PDIN (protein digestible in the small intestine, g/kg DM, when degradable nitrogen limits microbiological growth; INRA 2007) and PDIE (protein digestible in the small intestine, g/kg DM, when available energy limits microbial growth) are used. PDIN is the protein supplied by rumen-undegradable protein (PDIA) plus that supplied by microbial protein from rumen-degradable dietary protein. In comparison, PDIE is PDIA plus the microbial protein from rumen-fermented organic matter (INRA, 2007). At the end of the pre-experimental phase (0-10 d), heifers were group-housed indoors on deep straw bedding. They were fed a reconstituted milk replacer (MR) made from 135 g milk powder (23.9% crude protein and 19.0% fat content) and 865 g water per L until weaning (*ca*. 77-84 d of age). They were reared in dynamic groups: calves entered the group each week, while others left it at weaning. They were individually fed with automatic milk feeding systems (AMFS), with *ad libitum* access to fresh water, straw and hay. Group size ranged from 8-24 calves per AMFS. From day 11, milk was distributed according to the standard ration routinely used in the experimental herd (SD) or the standard ration increased by 15% (ID1 & ID2). All calves were fed TMR no. 1 (TMR1) *ad libitum* (Table 1).The TMR1 contained 47.5% of maize silage, 47.5%of concentrate 1 and 5% of 18 % CP lucerne pellets.

**Table 1.**
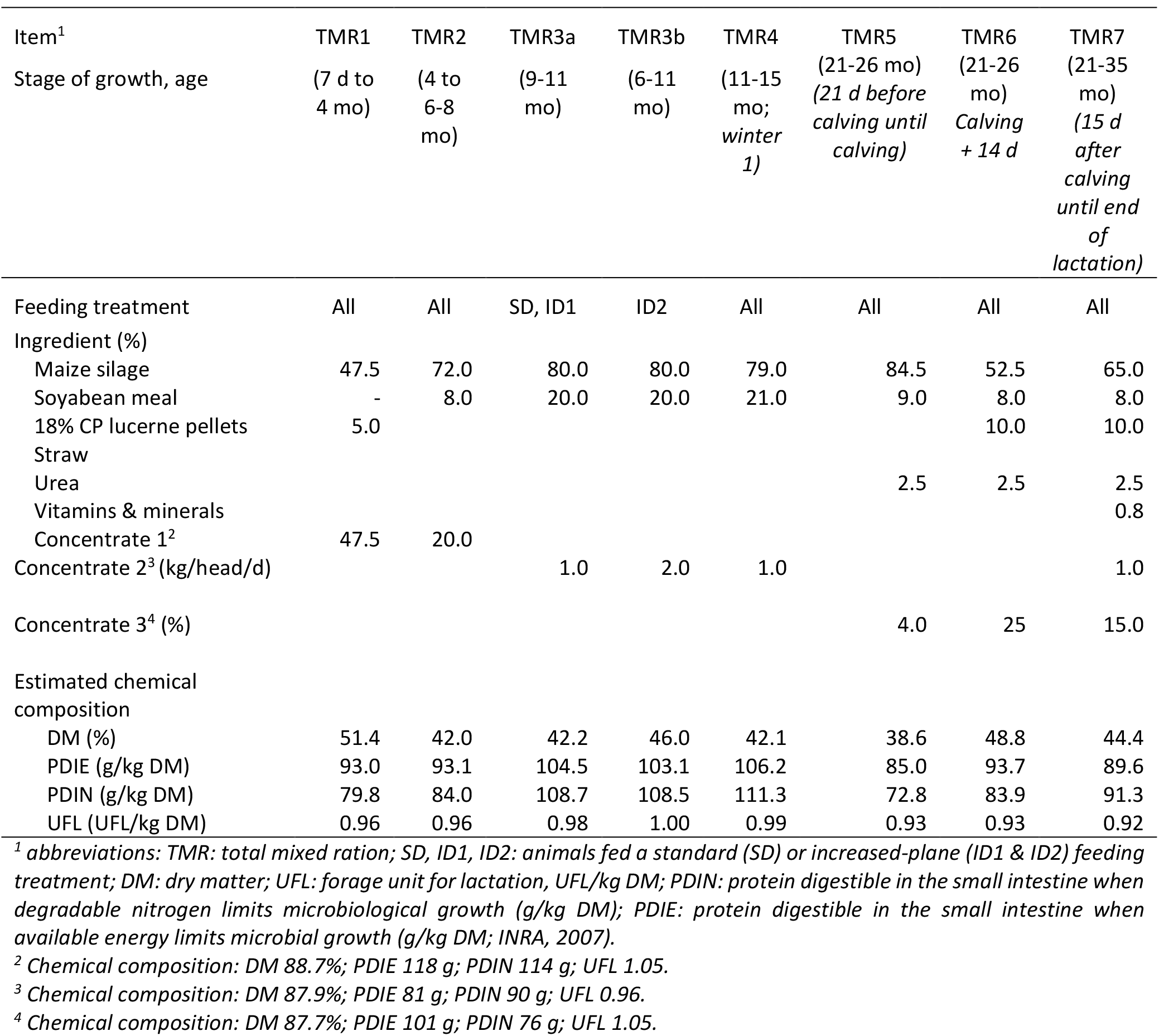
Ingredients and chemical composition of the experimental diets

From weaning to 6-8 mo of age, calves were housed on deep straw bedding with *ad libitum* access to fresh water and straw. Until 4 mo of age, the SD group received TMR1 *ad libitum* until the maximum daily allowance of concentrate intake reached 2 kg DM/head/d. No restriction was applied for ID1 or ID2 heifers. From 4 to 6-8 mo of age, TMR2 was distributed *ad libitum* until concentrate intake reached 2.0, 2.5 and 2.5 kg DM/head/d for SD, ID1 and ID2 heifers, respectively *i.e*. total daily allowance of 10.0, 12.5 and 12.5 kg DM/head/d, respectively. These amounts did not change until being turned out to pasture. The TMR2 contained 72% of maize silage, 8% of soya bean meal and 20% of concentrate 1.

SD, ID1 and ID2 heifers were turned out to pasture from mid-May, mid-May and mid-June, respectively, and rotationally grazed on a perennial ryegrass sward. After a 5-d transition phase and throughout the grazing season, the SD and ID1 groups received a supplement of 1 kg DM/heifer/d of concentrate 2. The ID2 group received 1 kg DM/heifer/d of maize silage and 2 kg DM/heifer/d of concentrate 2. Grass availability and/or quality were insufficient to maintain the desired growth rates during summer. SD and ID1 heifers then received up to 2.5 kg DM/heifer/d of additional TMR3a, plus 1 kg DM/heifer/d of concentrate 2. ID2 heifers received up to 3 kg DM/heifer/d of TMR3b, plus 2 kg DM/heifer/d of concentrate 2. To reach 380 kg at the end of the grazing season (when oestrus synchronisation started), the expected ADG for SD and ID1 heifers was *ca*. 600 g/d during this period, with a feeding regime based on grass plus 1 kg DM/heifer/d of concentrate 2, and 800 g/d when receiving grass plus TMR3a. For ID2 heifers, it was estimated that grass alone was not sufficient to reach 900 g/d during the same period, so TMR3b was used (Table 1). In the pasture area, a permanent headlock barrier (80 places on a concrete floor) was used daily to feed concentrate to SD and ID1 heifers. Heifers were locked in for 1 hour while eating to decrease competition between heifers for feed. Since the ID2 group had *ad libitum* access to the ration, its heifers were not locked in. At the end of the first grazing season (the first week of November), heifers were group-housed (8 heifers/pen) on deep straw bedding and received 3.8 kg DM/head/d of a diet containing 79% maize silage and 21% soya bean meal. They had *ad libitum* access to fresh water, straw and mineral supplements. Vitamins and minerals, when not included in the concentrate during rearing, were included in mineral blocks that contained 2.5% Ca, 2.0% Mg and 32.5% Na per kg of DM, as well as (in mg/kg) Zn (10 000), Mn (8250), Cu (1500), I (200), Se (20) and Co (13). The concentrates during growth contained 4% P, 27% Ca, 5% Mg, plus vitamins (in UI/kg; 1 000 000 vitamin A, 350 000 vitamin D3 and 8 000 vitamin E). They also contained (in mg/kg) Cu (1500), Zn (10 000), I (200), Co (100) and Se (10). During lactation, the mineral supplement contained 7% P, 22% Ca and 4% Mg, plus vitamins (in UI/kg; 500 000 vitamin A, 100 000 vitamin D3 and 1 500 vitamin E). It also contained (in mg/kg) Cu (1000), Mn (3500), Zn (4530), I (80), Co (35) and Se (22).

After a 2-week adaptation period, heifers’ oestrous cycles were synchronised (see below), and the same rearing procedure was applied to all heifers. Heifers were turned out to pasture (generally in March) based on the date of successful insemination. They were reared in a single group and received no additional feed except for grass, along with the supplemental vitamins and minerals.

All heifers were housed indoors three weeks before the expected date of calving, along with multiparous cows, in a cubicle barn with fresh straw bedding that was distributed daily. Heifers were fed individually and received TMR5 daily, composed of maize silage (84.5%), soya bean meal (9%), concentrate (4%) and straw. From calving to 14 d post-calving, cows individually received TMR6, which contained maize silage (52.5%), soya bean meal (8%), concentrate (25%), dehydrated lucerne (1%), vitamin/mineral supplements, urea and straw (Table 1).

From day 14 after calving, cows individually received TMR7, which contained maize silage (65%), soya bean meal (8%), concentrate (15%), dehydrated lucerne (1%), urea and vitamin/mineral supplements (7% P, 22% Ca and 4% Mg). All cows were fed *ad libitum* during lactation assuming at least 10% refusal per day. Feed was distributed twice per day (08:00 and 17:00), and refusals were collected each morning (7:00) before fresh TMR was distributed.

The chemical composition of TMR ingredients produced on-farm (maize silage, straw) was determined at harvest, and an average sample of each, came from daily sample, was analysed. Another analyse was also done when the storage silo of maize silage changed. However, DM was determined at least once a week for all TMR ingredients. A similar procedure was applied to concentrate feed. The manufacturer analysed the feed (*e.g*. concentrate, soya bean) before delivering it, and we compared it to the average sample when changing feed. The estimated chemical composition of TMR was then determined using INRAtion^®^ software (INRA, 2010) based on these analyses and the percentage of each ingredient in the TMR. Due to potential changes in composition (*e.g*. DM or grain content of maize silage), TMR composition was checked regularly, and the amount of each ingredient was adapted accordingly. Grass intake was not measured. All heifers and cows housed indoors had *ad libitum* access to fresh water during the entire experiment.

### Age at first service

Age at first service (AFS) was then classified to understand better which factors could influence AFS and how future performance may be related to AFS. Three classes were created, with nearly an equal number of animals in each (Table 2).

**Table 2.** Description of the classes of age (in mo) at first service (AFS)

| Characteristic | AFS<sub>12.5</sub> | AFS<sub>14.0</sub> | AFS<sub>15.5</sub> |
| --- | --- | --- | --- |
| AFS<sup>1</sup> | 12.6 (0.73) | 14.2 (0.36) | 15.4 (0.65) |
| Total heifers | 58 | 57 | 60 |
| Heifers in SD | 16 | 29 | 29 |
| Heifers in ID1 | 15 | 27 | 30 |
| Heifers in ID2 | 27 | 1 | 1 |

<sup>1</sup>Mean (and standard deviation) of age at first service (AFS)

### Oestrus synchronisation

All heifers were inseminated after oestrus synchronisation during the second winter of rearing so that calving would occur at *ca*. 24 mo of age. At the end of November, oestrus was synchronised for nearly half of the heifers using a progestin ear implant (Norgestomet®, Intervet, Angers, France) along with an intramuscular injection of oestrogen (Crestar®, Intervet, Angers, France), without considering ovarian activity. A second synchronisation was performed three weeks later for the remaining heifers. The ear implant was removed after 9 d of treatment. Heifers generally showed signs of oestrus within 24-96 h and were inseminated when oestrus was detected. Heifers that failed to conceive but exhibited further signs of oestrus were inseminated at the end of the reproductive season (April). Ultrasonography was conducted an average of 42 d after insemination to determine pregnancy. Non-gestating heifers were excluded from the rest of the experiment.

### Sampling and measurements

Heifers were weighed every 14 d from birth to weaning, every 21 d from weaning until being turned out to pasture and every 28 d until the end of the experiment. BW was interpolated to compare the BW of heifers at similar stages of growth. ADGs were then calculated. Heifer health and care information was recorded throughout the experiment. The body condition score (BCS) was recorded three weeks before the expected date of calving and then once a month. The method and scale (ranging from 0-5) developed by Bazin *et al*. (1984) was used. BCS was scored by three trained technicians, whose scores were averaged.

Five measurements were recorded to monitor morphological traits during rearing and first lactation: heart girth (HG), chest depth, wither height (WH), hip width and backside width. A tape measure was used to measure HG, while a height gauge was used for the other measurements. The measurements were recorded only for the two first cohorts (2009-10 and 2010-11: Appendix Fig.1). Results were interpreted by class of age at first service calving (AFC), which was created later (not shown or discussed in the present article).

Daily feed intake was calculated individually as the daily feed allowance minus refusals. The allowance and refusals were assumed to have the same composition. DM of silage was determined five times per week, while DM of the pellets was determined once per week. Feed composition was estimated from average samples for maize silage, straw, soya bean and concentrate. Composition was not available for fresh grass (Table 1).

### Milk content analysis

Milk yield was automatically recorded at each milking (*i.e*. twice per day). During six successive milkings (Tuesday-Thursday), milk samples were collected and analysed for each cow to determine the fat and protein contents (Milkoscan, Foss Electric, Hillerod, Denmark). Fat- and-protein-corrected milk (FPCM, kg) was calculated using the following equation (INRA, 2018):

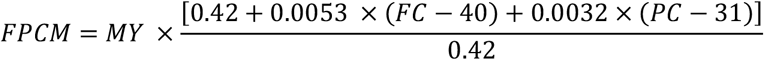

where FC is milk fat content (g/kg), PC is milk protein content (g/kg) and 0.42 is the UFL value for 1 kg of milk containing 40 g/kg of fat and 31 g/kg of protein.

### Milk progesterone analysis

Morning milk samples were collected Monday, Wednesday and Friday from calving to two weeks after the service that induced pregnancy, or five weeks after the end of the breeding season (*i.e*. July), and were then stored at −20°C to determine progesterone using commercial ELISA kits (Milk Progesterone ELISA, Ridgeway Science Ltd., England). Coefficients of variation among assays for ELISA on 5 ng/ml control samples ranged from 8-14% among experimental years.

#### Determining Luteal Activity

Two progesterone (P4) milk concentration thresholds were defined, following Petersson *et al*. (2006) and adapted by Cutullic *et al*. (2011), to distinguish (i) the baseline P4 level in milk from the luteal phase level (threshold 1) and (ii) a low luteal phase level from a high luteal phase level (threshold 2). P4 values were classified as negative (< threshold 1), positive (> threshold 2) or intermediate. An increase in P4 milk concentrations was considered to be induced by corpus luteum activity when at least two consecutive values were not negative and at least one was positive. Due to the sampling schedule (Monday, Wednesday and Friday), the interval between samples was 2 d or 3 d. A decrease in P4 milk concentration was considered to result from luteolysis of the corpus luteum when at least one value became negative. These definitions helped to identify and distinguish luteal phases from inter-luteal phases.

#### Qualifying Progesterone Profiles

Physiological intervals were calculated for each luteal phase: commencement of luteal activity (CLA), cycle length (IOI), luteal phase length (LUT) and inter-luteal interval (ILI; for details, see Cutullic *et al*., 2011). Ovulation was considered to induce a prolonged luteal phase (PLP) if the luteal phase exceeded 25 d. Ovulation was considered to be delayed if the inter-luteal interval exceeded 12 d. Based on these definitions, P4 profiles were classified as (i) normal, (ii) PLP profile (when at least one PLP was observed), (iii) delayed (D; if CLA > 60 d), (iv) interrupted (I; when at least one ovulation > 2 was delayed) and (v) disordered (Z; when luteal activity appeared irregular but could not be assigned to another abnormality class).

### Calculations and statistical analysis

All data on dairy cows (*e.g*. reproduction, milk yield, feed intake) was automatically stored in a dedicated recording system. Analyses of heifer growth and performance, as well as data on progesterone, were recorded in Microsoft Excel files. All data manipulation and statistical analyses were performed in R software using the *lm* procedure for ANOVA or *glm* for logistic regressions (R Core Team, 2019). Normal distribution of the residuals, equality of the variance and non-dependent data were checked for all models. Quantitative traits (*i.e*. growth, age, BW, milk yield, BCS, CLA, cycle lengths) were studied using the following ANOVA model:

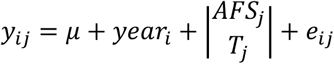

where y_ij_ is the variable of interest, μ is the overall mean of the variable of interest, year_i_ is the fixed effect of the experimental year (i=1, 2 or 3), AFS_j_ is the fixed effect of AFS (j= 12.5, 14.0 or 15.5 mo) or T_j_ is the fixed effect of feeding treatment (j= SD, ID1 or ID2) included in the model, and e_ij_ is the random residual effect. Year was included as a fixed effect because there were only three levels (year1, year2, year3), and this approach seemed the most appropriate option given the small number of levels. Had year been included as a random effect, variance would have been estimated from only three levels, rendering it inaccurate.

Dichotomous traits (*i.e*. reproductive success and type of cyclicity pattern) were studied using the following logistic regression model:

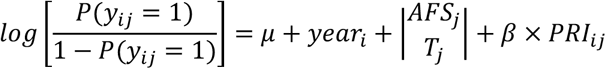

where y_ij_ is the variable of interest, μ is the overall mean and the fixed effects (yeari, AFS_j_ or T_j_) are the same as previously described.

For the reproductive performance of heifers, the covariate PRI_ij_ was added; it describes the effect of the interval from the removal of the last progesterone-releasing implant until insemination. This covariate was not required for the performance of cows because only heifers were synchronised. Effects were considered highly significant at P<0.001, significant at P<0.05 and a trend at P<0.10.

## Results

Of the 217 heifers in the experiment, 175 successfully calved. The 42 that did not either died during rearing (7), were culled due to injuries (6) or were not pregnant within the breeding period considered for the present study (29).

### Growth and reproductive performance of heifers

Mean BW at birth was 41.3 kg (± 5.2) and did not differ significantly among all groups (*i.e*. not associated with the feeding treatment, P = 0.85, or AFS, P = 0.15; Table 3; Table 4).

**Table 3.** Effects of feeding treatment on the growth and reproductive performance of heifers during the rearing period

|  | Feeding Treatment |  |  | Model<sup>1</sup> |  | Significance levels<sup>2</sup> |
| --- | --- | --- | --- | --- | --- | --- |
|  | SD1 | ID1 | ID2 | R<sup>2</sup><sub>adj</sub> | RSE |  |
| Number of heifers | 74 | 72 | 29 |  |  |  |
| **Growth** |  |  |  |  |  |  |
| BW<sup>3</sup> at birth (kg) | 41.2 | 41.7 | 41.1 | 0.00 | 5.19 | 0.85 |
| BW at first AI (kg) | 400.7<sup>a</sup> | 398.5<sup>a</sup> | 378.1<sup>b</sup> | 0.14 | 33.29 | ★★ |
| ADG<sup>4</sup> 0-6 months (g/d) | 958<sup>a</sup> | 976<sup>a</sup> | 1042<sup>b</sup> | 0.09 | 97.7 | ★★★ |
| ADG 6-12 months (g/d) | 703<sup>a</sup> | 699<sup>a</sup> | 789<sup>b</sup> | 0.31 | 116.8 | ★★ |
| ADG 12-18 months (g/d) | 800<sup>a</sup> | 774<sup>a</sup> | 660<sup>b</sup> | 0.11 | 133.2 | ★★★ |
| **Reproduction** |  |  |  |  |  |  |
| Start of breeding season to first service interval (d) | 13.9 | 12.8 | 14.0 | 0.00 | 5.76 | 0.46 |
| Pregnancy rate at first service (%) | 64 | 58 | 66 | NA | NA | 0.64 |
| Number of services | 1.9 | 1.8 | 1.5 | 0.21 | 0.78 | ● |
| Pregnant (%) | 95 | 96 | 90 | NA | NA | 0.67 |
| Age at first calving (months) | 24.0<sup>a</sup> | 23.9<sup>a</sup> | 21.9<sup>b</sup> | 0.32 | 1.26 | ★★★ |
| Calf BW (kg) | 38.4 | 37.6 | 37.2 | 0.32 | 4.02 | 0.37 |

<sup>1</sup>adjusted coefficient of determination: R<sup>2</sup><sub>adj</sub>; residual standard error: RSE
<sup>2</sup> ★★★ $P < 0.001$ ; ★★ $P < 0.01$ ; ★ $P < 0.05$ ; ● $P < 0.1$ ; otherwise, the exact P-value
<sup>3</sup>body weight: BW
<sup>4</sup>average Daily Gain: ADG
<sup>a-b</sup> Different superscripts indicate adjusted means that differ between feeding treatments ( $P < 0.05$ , Tukey's pairwise comparison)

**Table 4.** Relations between age at first service and growth and reproductive performance of heifers during the rearing period

|  | Age at first service (AFS) |  |  | Model<sup>1</sup> |  | Significance levels<sup>2</sup> |
| --- | --- | --- | --- | --- | --- | --- |
|  | AFS<sub>12.5</sub> | AFS<sub>14.0</sub> | AFS<sub>15.5</sub> | R<sup>2</sup><sub>adj</sub> | RSE |  |
| Number of heifers | 58 | 57 | 60 |  |  |  |
| **Growth** |  |  |  |  |  |  |
| BW<sup>3</sup>at birth (kg) | 41.5 | 42.0 | 40.2 | 0.02 | 5.13 | 0.15 |
| BW at first AI<sup>4</sup> (kg) | 373.1<sup>a</sup> | 394.3<sup>b</sup> | 419.8<sup>c</sup> | 0.37 | 28.49 | ★★★ |
| ADG<sup>5</sup> 0-6 months (g/d) | 1001 | 960 | 978 | 0.03 | 100.8 | ● |
| ADG 6-12 months (g/d) | 759<sup>a</sup> | 688<sup>b</sup> | 698<sup>b</sup> | 0.30 | 117.5 | ★★ |
| ADG 12-18 months (g/d) | 712<sup>a</sup> | 799<sup>b</sup> | 790<sup>b</sup> | 0.07 | 136.3 | ★★ |
| **Reproduction** |  |  |  |  |  |  |
| Start of breeding season to first service interval (d) | 12.9 | 13.2 | 14.3 | 0.00 | 5.75 | 0.42 |
| Pregnancy rate at first service (%) | 59 | 60 | 67 | NA | NA | 0.30 |
| Number of services | 1.7 | 1.7 | 1.9 | 0.20 | 0.78 | 0.25 |
| Pregnant (%) | 93 | 91 | 98 | NA | NA | 0.37 |
| Age at first calving (months) | 22.3<sup>a</sup> | 23.8<sup>b</sup> | 24.8<sup>c</sup> | 0.52 | 1.06 | ★★★ |
| Calf body weight (kg) | 37.4 | 38.6 | 37.7 | 0.32 | 4.02 | 0.31 |

<sup>1</sup>adjusted coefficient of determination: R<sup>2</sup><sub>adj</sub>; residual standard error: RSE
<sup>2</sup>★★★ P < 0.001; ★★ P < 0.01; ★ P < 0.05; ● P < 0.1; otherwise, the exact P-value
<sup>3</sup>body weight: BW
<sup>4</sup>artificial insemination: IA
<sup>5</sup>average Daily Gain: ADG.
<sup>a-b</sup> Different superscripts indicate adjusted means that differ between feeding treatments (P < 0.05, Tukey's pairwise comparison)

The feeding treatment had little effect on growth during the milking phase, and heifers reached 117 kg (± 11.8) at 3 mo of age (immediately after weaning). From weaning to 6 mo, heifers in the ID2 treatment were heavier than those in the SD and ID1 treatments (229 kg *vs* 213 kg and 217 kg at 6 mo, respectively; P < 0.001; Fig. 1A). The highest ADG was observed for ID2 heifers from 0-6 mo (1042 *vs* 958 and 976 g/d for ID2, SD and ID1, respectively; P < 0.001, Table 3). This difference remained significant from 6-12 mo of age (789, 703 and 699 g/d for ID2, SD and ID1 heifers, respectively; P 0.01, Table 3). However, from 12-18 mo, ADG was significantly lower for ID2 heifers than for SD and ID1 heifers (660 *vs* 800 and 774 g/d, respectively; P < 0.001, Table 3).

**Figure 1.**
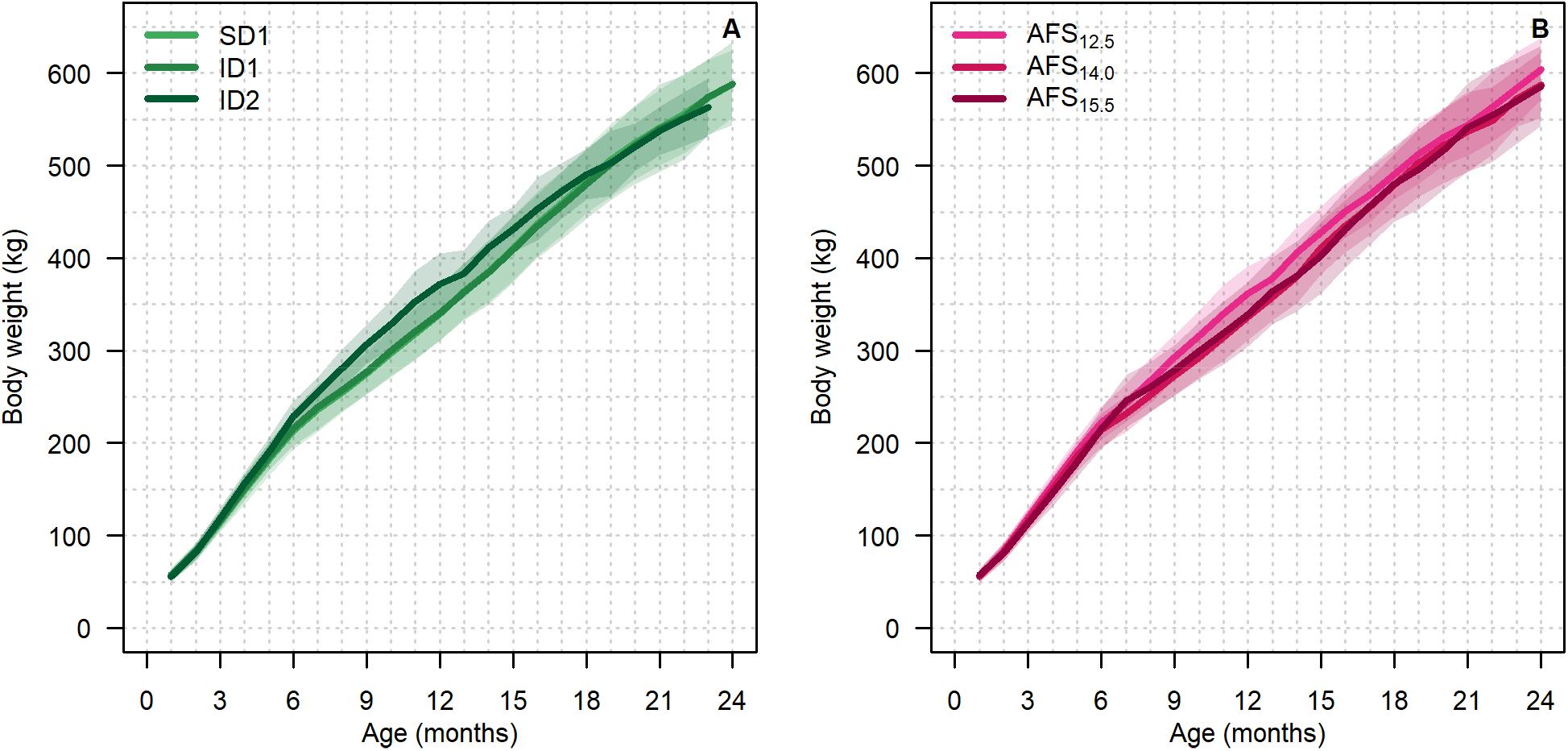
Mean body weight of heifers during the rearing period by (A) feeding treatment (SD, ID1, ID2: animals fed a standard (SD) or increased-plane (ID1 & ID2) feeding treatment) and (B) class of age at first service (AFS). Shaded areas are the dispersions of the data around the means (± one standard deviation).

The feeding treatment had no effect on reproductive performance (Table 3), although ID2 heifers tended to have fewer services than SD or ID1 heifers (1.5 vs 1.9 or 1.8, respectively). Cows in the three feeding treatments had a similar interval from the start of the breeding season to the first service (13.5 d), similar success at the first service (*ca*. 62% of heifers pregnant) and a similar pregnancy rate by the end of the breeding season (94%).

No difference in calf BW (37.9 kg) was observed, despite a difference in their dam’s BW at the first service and first calving (ID2 heifers were lighter than SD and ID1 heifers Table 3 and 5). ID2 heifers calved at a younger age than SD or ID1 heifers (*ca*. 2 mo earlier, P < 0.001; Table 3).

**Table 5.** Effects of feeding treatment during the rearing period on productive and reproductive performances of primiparous cows

|  | Feeding Treatment |  |  | Model<sup>1</sup> |  | Significance levels<sup>2</sup> |
| --- | --- | --- | --- | --- | --- | --- |
|  | SD1 | ID1 | ID2 | R<sup>2</sup><sub>adj</sub> | RSE |  |
| Number of cows | 67 | 68 | 24 |  |  |  |
| **Production** |  |  |  |  |  |  |
| Total milk yield per 308 d (kg) | 7312 | 7370 | 6920 | 0.19 | 706.9 | ● |
| Peak milk yield (kg) | 31.3<sup>a</sup> | 31.9<sup>a</sup> | 28.7<sup>b</sup> | 0.10 | 3.50 | ★★ |
| Mean fat content (g/kg) | 37.0 | 36.5 | 36.2 | 0.10 | 3.66 | 0.75 |
| Mean protein content (g/kg) | 30.2 | 29.7 | 29.4 | 0.02 | 1.53 | 0.17 |
| Fat- and protein- corrected milk (kg) | 6983<sup>a</sup> | 6973<sup>a</sup> | 6138<sup>b</sup> | 0.26 | 668.5 | ★ |
| **Conformation** |  |  |  |  |  |  |
| BW<sup>3</sup> at first calving (kg) | 542<sup>a</sup> | 534<sup>a</sup> | 501<sup>b</sup> | 0.10 | 43.0 | ★★★ |
| BCS<sup>4</sup> at calving (0-5 scale) | 2.45 | 2.40 | 2.30 | 0.33 | 0.296 | 0.11 |
| BCS at nadir (0-5 scale) | 1.85 | 1.80 | 1.75 | 0.43 | 0.267 | 0.47 |
| BCS loss to nadir (0-5 scale) | -0.55 | -0.60 | -0.60 | 0.44 | 0.255 | 0.81 |
| **Cyclicity<sup>5</sup>** |  |  |  |  |  |  |
| CLA (d) | 20.9 | 24.8 | 20.1 | 0.00 | 0.56 | 0.23 |
| IOI<sub>1</sub> (d) | 20.7 | 23.8 | 24.9 | 0.04 | 14.01 | 0.47 |
| LUT<sub>1</sub> (d) | 13.3 | 13.9 | 14.9 | 0.18 | 10.77 | 0.88 |
| ILI<sub>1</sub> (d) | 9.6 | 11.2 | 7.7 | 0.04 | 11.29 | 0.55 |
| IOI<sub>2-4</sub> (d) | 23.3 | 23.6 | 21.2 | 0.00 | 5.91 | 0.42 |
| LUT<sub>2-4</sub> (d) | 13.8 | 13.7 | 12.5 | 0.39 | 5.79 | 0.77 |
| ILI<sub>2-4</sub> (d) | 9.0 | 10.2 | 9.0 | 0.45 | 4.76 | 0.54 |
| Normal (%) | 65% | 59% | 53% | NA | NA | 0.52 |
| PLP (%) | 19% | 18% | 33% | NA | NA | 0.44 |
| Delayed (%) | 10% | 12% | 7% | NA | NA | 0.81 |
| **Fertility** |  |  |  |  |  |  |
| Number of services per cow | 1.9<sup>a</sup> | 2.4<sup>b</sup> | 2.2<sup>ab</sup> | 0.10 | 1.27 | ★ |
| Pregnant (%) | 86% | 85% | 87% | NA | NA | 0.92 |
| Calf BW (kg) | 38.4 | 37.8 | 36.9 | 0.00 | 4.84 | 0.40 |

<sup>1</sup>adjusted coefficient of determination: R<sup>2</sup><sub>adj</sub>; residual standard error: RSE
<sup>2</sup>★★★ P < 0.001; ★★ P < 0.01; ★ P < 0.05; ● P < 0.1; otherwise, the exact P-value
<sup>3</sup>body weight: BW
<sup>4</sup>body condition score: BCS
<sup>5</sup>commencement of luteal activity: CLA; cycle length: IOI; luteal phase length : LUT; inter-luteal interval : ILI; prolonged luteal phase: PLP
<sup>a-b</sup> Different superscripts indicate adjusted means that differ between feeding treatments (P < 0.05, Tukey's pairwise comparison)

Heifers inseminated at the youngest age (a mean of 12.5 mo; AFS_12.5_) tended to have a higher growth rate from 0-6 mo of age than those inseminated at 14.0 (AFS_14.0_) or 15.5 (AFS_15.5_) mo of age (1001 *vs* 960 or 978 g/d, respectively; P < 0.10; Table 4). This difference increased from 6-12 mo of age (759 *vs* 688 and 698 for AFS_12.5_, AFS_14.0_ and AFS_15.5_, respectively; P < 0.01; Table 4; Fig. 1B).

From 12-18 mo of age, AFS_12.5_ heifers had a lower growth rate than AFS_14.0_ and AFS_15.5_ heifers ADG of 712 *vs* 799 and 790 g/d, respectively (P < 0.001; Table 4). This is consistent with the effects of the feeding treatment and the distribution of animals among the AFS classes and feeding treatments (Table 2).

AFS had no influence on fertility (Table 4). All heifers had a similar interval from the start of the breeding season to the first service, a similar success at the first service and a similar pregnancy rate by the end of the breeding season, with a similar number of services per animal.

No difference in calf BW (37.9 kg) was observed, despite a difference in the dam’s BW at first service and at first calving (AFS_12.5_ heifers were lighter than those in AFS_14.0_, which were lighter than those in AFS_15.5_, Tables 4 and 6). Consistent with the AFS, AFS_12.5_ heifers calved younger than AFS_14.0_ heifers, which calved younger than AFS_15.5_ heifers (Table 4).

**Table 6.** Effects of the class of age at first service (AFS) on the productive and reproductive performance of primiparous cows

|  | Age at first service (AFS) |  |  | Model<sup>1</sup> |  | Significance levels<sup>2</sup> |
| --- | --- | --- | --- | --- | --- | --- |
|  | AFS<sub>12.5</sub> | AFS<sub>14.0</sub> | AFS<sub>15.5</sub> | R<sup>2</sup><sub>adj</sub> | RSE |  |
| Number of cows | 51 | 50 | 58 |  |  |  |
| **Production** |  |  |  |  |  |  |
| Total milk yield per 308 d (kg) | 7229 | 7236 | 7370 | 0.15 | 721.7 | 0.68 |
| Peak milk yield (kg) | 30.2 | 31.6 | 31.7 | 0.04 | 3.59 | ● |
| Mean fat content (g/kg) | 36.2 | 36.9 | 36.8 | 0.10 | 3.65 | 0.66 |
| Mean protein content (g/kg) | 29.8 | 29.9 | 29.9 | 0.00 | 1.56 | 0.93 |
| Fat- and protein- corrected milk (kg) | 6800 | 6891 | 7000 | 0.26 | 688.4 | 0.51 |
| **Conformation** |  |  |  |  |  |  |
| BW<sup>3</sup> at first calving (kg) | 509<sup>a</sup> | 539<sup>b</sup> | 549<sup>b</sup> | 0.14 | 41.9 | ★★★ |
| BCS<sup>4</sup> at calving (0-5 scale) | 2.35<sup>a</sup> | 2.35<sup>a</sup> | 2.45<sup>b</sup> | 0.34 | 0.295 | 0.05 |
| BCS at nadir (0-5 scale) | 1.75 | 1.8 | 1.85 | 0.44 | 0.264 | 0.13 |
| BCS loss to nadir (0-5 scale) | -0.60 | -0.60 | -0.55 | 0.44 | 0.254 | 0.41 |
| **Cyclicity<sup>5</sup>** |  |  |  |  |  |  |
| CLA (d) | 20.2 | 23.6 | 23.7 | 0.00 | 0.56 | 0.39 |
| IOI<sub>1</sub> (d) | 25.0 | 19.8 | 23.2 | 0.04 | 13.96 | 0.31 |
| LUT<sub>1</sub> (d) | 13.9 | 12.3 | 14.9 | 0.19 | 10.73 | 0.57 |
| ILI<sub>1</sub> (d) | 10.7 | 8.7 | 10.7 | 0.04 | 11.32 | 0.68 |
| IOI<sub>2-4</sub> (d) | 23.0 | 22.3 | 24.1 | 0.00 | 5.92 | 0.45 |
| LUT<sub>2-4</sub> (d) | 14.5 | 13.6 | 12.7 | 0.39 | 5.75 | 0.44 |
| ILI<sub>2-4</sub> (d) | 8.8 | 8.8 | 11.1 | 0.48 | 4.67 | ● |
| Normal (%) | 58 | 68 | 56 | NA | NA | 0.55 |
| PLP (%) | 29 | 8 | 23 | NA | NA | ★ |
| Delayed (%) | 5 | 13 | 14 | NA | NA | 0.23 |
| **Fertility** |  |  |  |  |  |  |
| Number of services per cow | 1.9 | 2.4 | 2.2 | 0.08 | 1.28 | 0.16 |
| Pregnant (%) | 86% | 88% | 84% | NA | NA | 0.90 |
| Calf BW (kg) | 37.2<sup>a</sup> | 39.3<sup>b</sup> | 37.3<sup>a</sup> | 0.04 | 4.77 | ★ |

<sup>1</sup>adjusted coefficient of determination: R<sup>2</sup><sub>adj</sub>; residual standard error: RSE
<sup>2</sup>★★★ P < 0.001; ★★ P < 0.01; ★ P < 0.05; ● P < 0.1; otherwise, the exact P-value
<sup>3</sup>body weight: BW
<sup>4</sup>body condition score: BCS
<sup>5</sup>commencement of luteal activity: CLA; cycle length: IOI; luteal phase length : LUT; inter-luteal interval : ILI; prolonged luteal phase: PLP
<sup>a-b</sup> Different superscripts indicate adjusted means that differ between feeding treatments (P < 0.05, Tukey's pairwise comparison)

### Lactating performance of primiparous cows

BW recorded immediately after calving was lower for ID2 cows than for SD and ID1 cows (501 *vs* 542 and 534 kg, respectively; P < 0.001; Table 5; Fig. 2A.), which is consistent with the observation that ID2 heifers first calved younger than SD and ID1 heifers (Table 4). No difference in BCS was observed among the feeding treatments during the first lactation (result not shown). On a 308 d basis, ID2 cows tended to produce less milk than SD and ID1 cows (6920 *vs* 7312 and 7370 kg, respectively; P < 0.10; Table 5; Fig. 2C). No difference in mean fat and protein contents was observed among feeding treatments. However, cows that received the ID2 treatment when heifers produced less FPCM than cows that received the SD or ID1 treatments (6482 *vs* 6983 and 6973 kg, respectively; P < 0.05). ID2 cows had a lower peak milk yield than SD and ID1 cows (28.7 *vs* 31.3 and 31.9 kg/d, respectively; P < 0.001). During the first seven weeks of lactation, ID2 cows were lighter (on average, 38 and 25 kg less than SD and ID1 cows, respectively), and produced less milk (3.1 kg/d less than SD and ID1). This difference decreased during the last part of the period (8-15 weeks); ID2 cows weighed 27 and 17 kg less than SD and ID1 cows, respectively, and produced 2.2 and 2.9 kg/d less milk than SD and ID1 cows, respectively.

**Figure 2.**
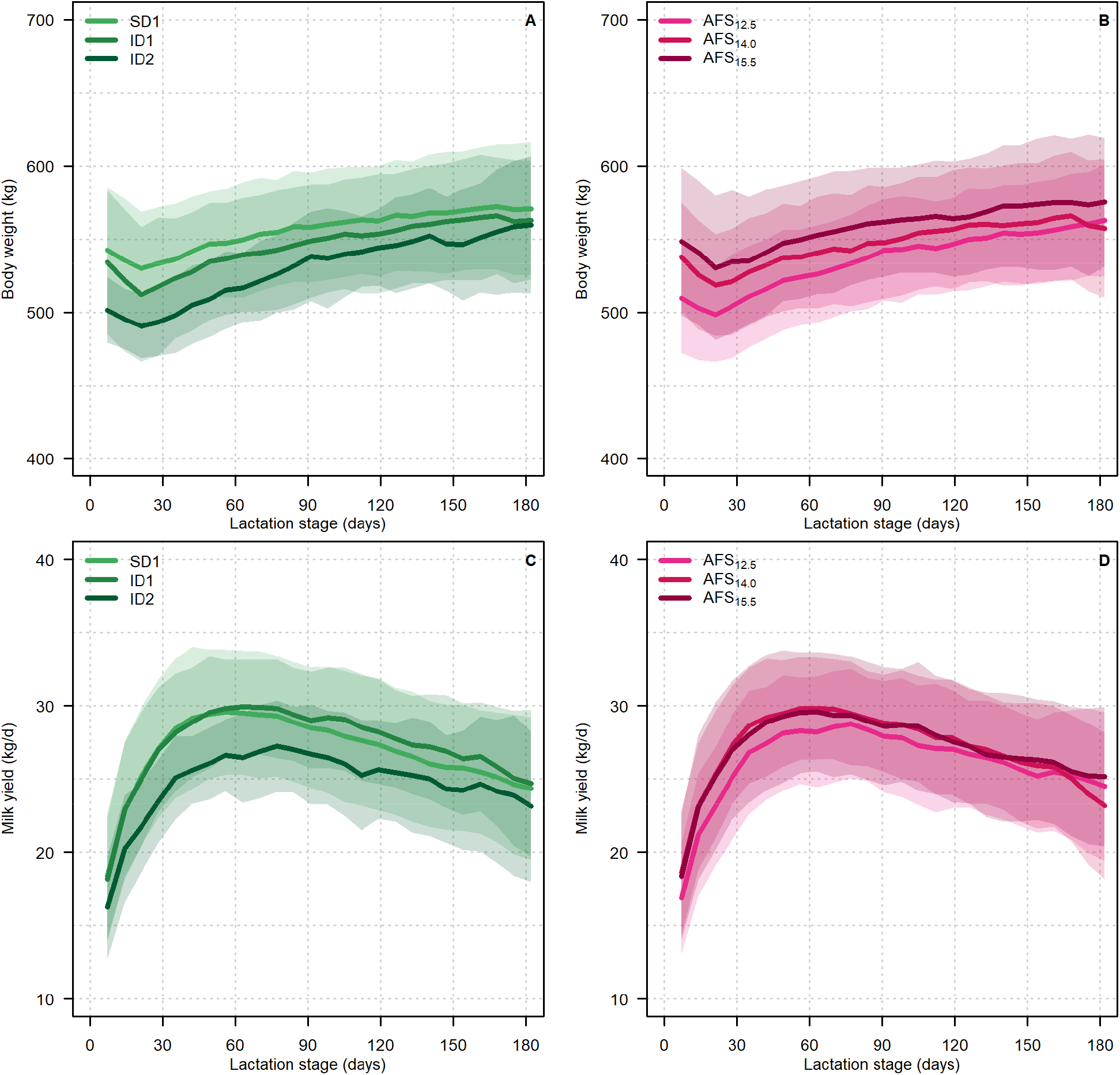
(A and B) body weight and (C and D) milk yield of primiparous cows during lactation by (A and C) feeding treatment (SD, ID1, ID2: animals fed a standard (SD) or increased-plane (ID1 & ID2) feeding treatment) and (B and D) class of age at first service (AFS). Shaded areas are the dispersions of the data around the means (± one standard deviation).

The feeding treatment of dairy cows during the rearing period did not affect ovarian cyclicity during the first lactation (Table 5). Mean CLA was 20.4 d, and the first IOI was 20.7 d, with no difference in LUT or ILI among treatments. No difference in the subsequent cycles was observed, with a mean IOI of 23.3 d. The distribution of abnormal patterns of ovarian activity was not significant, although ID2 cows had a lower normal profile rate than ID1 cows, which had a lower normal profile rate than SD cows (53% *vs* 59% *vs* 65%, respectively; Table 5).

ID2 cows had an incidence of PLP abnormalities of 33%, while that for ID1 and SD cows was 18% and 19%, respectively (Table 5). *Ca*. 86% of cows were pregnant at the end of the breeding season, which had no relationship with feeding treatment. Although the difference in cyclicity among feeding treatments did not influence the re-calving rate, ID1 cows required more services for pregnancy to occur than SD cows (2.4 *vs* 1.9, respectively; P < 0.05; Table 5). The number of services required to achieve pregnancy was *ca*. 2.2 for ID2 cows. Feeding treatment had no influence on subsequent calf BW.

AFS influenced BW at calving, and was lower for AFS_12.5_ than for AFS_14.0_ and AFS_15.5_ cows (509 *vs* 539 and 549 kg, respectively, P < 0.001; Table 6; Fig. 2B). BCS at calving was higher for AFS_15.5_ than for AFS_12.5_ and AFS_14.0_ cows (2.45 *vs* 2.35 and 2.35, respectively; P < 0.05). After calving, BCS did not differ between AFS classes. On a 308 d basis, no difference in milk yield, composition or FPCM was observed. Only peak milk yield tended to be lower for AFS_12.5_ cows (30.2 kg) than for AFS_14.0_ and AFS_15.5_ cows (31.6 and 31.7 kg, respectively; Fig. 2D; Table 6).

AFS influenced fertility characteristics little. For ovarian cyclicity, all three AFS classes had similar CLA, with similar cycle lengths, except for AFS_15.5_ cows, which tended to have longer ILI from the second to fourth cycle than AFS_12.5_ and AFS_14.0_ cows (Table 6). AFS_14.0_ cows had a lower incidence of PLP than AFS_12.5_ and AFS_15.5_ cows (8% *vs* 29% and 23%, respectively; P < 0.05; Table 6). AFS did not influence fertility: all classes had similar number of services (2.2, on average), and an average of 86% of the cows in each class were pregnant at the end of the breeding season. Subsequent calf BW was heavier for AFS_14.0_ cows than for AFS_12.5_ and AFS_15.5_ cows (+2 kg; P < 0.05; Table 6). Feed intake did not differ among feeding treatments or among AFS classes (17 kg DM/d), even when it was corrected per kg of BW (Fig. 3).

**Figure 3.**
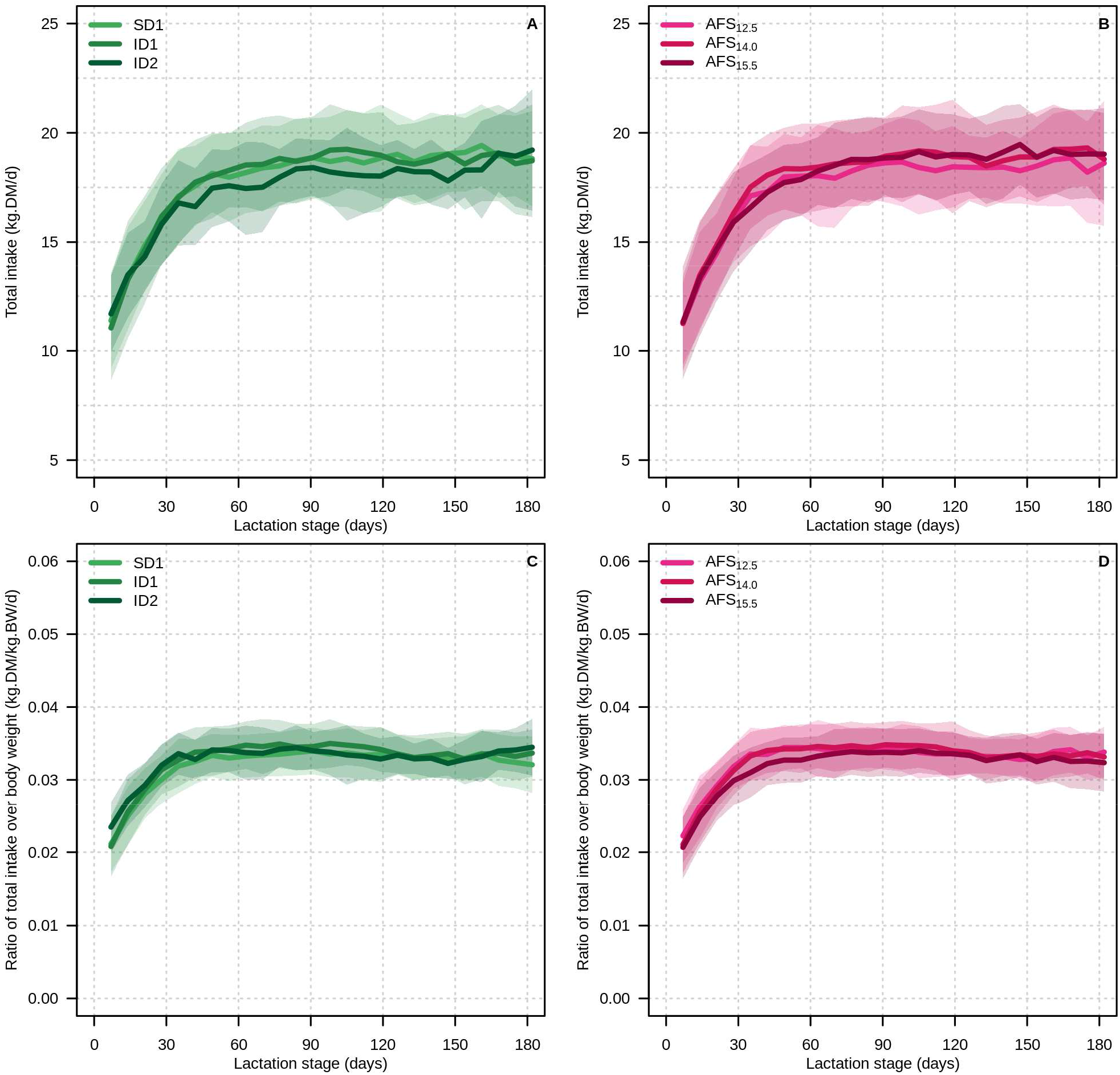
(A and B) daily dry matter intake and (C and D) daily ratio of dry matter intake over body weight of primiparous cows during lactation by (A and C) feeding treatment (SD, ID1, ID2: animals fed a standard (SD) or increased-plane (ID1 & ID2) feeding treatment) and (B and D) class of age at first service (AFS). Shaded areas are the dispersions of the data around the means (± one standard deviation).

Morphological trait analysis based on age at first calving (AFC) cohorts 2009-10 and 2010-11 (Appendix Fig.1) indicated that young cows at first calving (mean age of 21 mo, n = 30; AFC_21_) were lighter than those that first calved at a mean age of 23.5 mo (n = 39; AFC_23.5_) or 25 mo (n = 36; AFC_25_; 498 *vs* 528 and 563 kg, respectively; P < 0.05) and also had smaller morphological traits. For example, WH was 137.4, 139.1 and 140.4 cm for AFC_21_, AFC_23.5_ and AFC_25_, respectively; P < 0.05). However, at a given age (*e.g*. 25 mo), no difference among the three AFC treatments was observed (140.7, 140.4 and 142.0 mm, respectively).

## Discussion

The present study indicates that reducing the age of first service to *ca*. 12 mo and, consequently, age at first calving to 22 mo or less, influenced the performance of primiparous Holstein cows little. Several authors have shown that setting age at first calving of heifers at 23-26 mo of age increases longevity and maximises economic returns (Bach 2011; Wathes *et al*., 2014; Boulton *et al*., 2017). The early rearing period is key to reaching this target, as sub-optimal nutrition delays the onset of puberty, adversely affects skeletal growth and increases the risk of dystocia at first calving (Ettema and Santos 2004). Poor growth is the main reason for culling heifers prior to calving (Esslemont and Kossaibati 1997). Pre-weaning growth in dairy heifers is generally associated with the performance of first lactation (Khan *et al*. 2011; Soberon *et al*., 2012). Some studies reported that pre-weaning differences caused by different feeding regimes were not statistically significant as calves aged (Morrison *et al*. 2009; Quigley *et al*. 2006). This may be explained in part by a compensatory increase in growth when the feed allowance (*e.g*. level, energy, protein) is no longer limited after a period of restriction.

The differences in feed allowance resulted in differences in development and size at 6 and 12 mo of age but had little effect on BW at weaning. In a study by Johnson et al (2019), two treatment groups before weaning had significant differences in pre-weaning performance that persisted up to 6 mo. In our study, the high feed allowance before weaning, without restricting the TMR for control heifers, probably explains the lack of difference in BW observed at weaning. According to Morrison et al. (2009), on most commercial farms, a small amount of milk (4-6 L/day of whole milk or 400-600 g of milk replacer (MR) is offered until weaning at 42-56 days of age. According to Jasper and Weary (2002), *ad libitum* milk intake is *ca*. 12 L/day of whole milk, and intake in the present study was *ca*. 9 L/d per heifer until 11 weeks of age. The development and BW of animals at 6 mo were high (*e.g*. 111 cm HG and 220 kg BW), which fits well with recommendations for an optimal age at first calving at 24 mo of age or less. In a study by Ettema and Santos (2004) on the importance of age and BW at first calving for Holstein heifers, only 2.7% of dairy farms reached the recommended target BW, which resulted in economic losses. Total nutrient intake, energy source and protein content in the diet have a cumulative effect on how calves partition nutrients into tissue (Van Amburgh and Drackley 2005). During the milking phase, calves benefit when MRs contain more protein and less fat, and reach higher levels of skeletal growth (Hill *et al*., 2010). Therefore, providing more MR improves growth and feed efficiency (Bartlett *et al*., 2006). Increased nutrient intake is also associated with increased plasma levels of insulin-like growth factor 1 (Smith *et al*., 2002; Bartlett *et al*., 2006), which in part regulates the subsequent growth rate (Hammon *et al*., 2002; Brickell *et al*., 2009a). Several studies discuss effects of intensive growth during rearing (Le Cozler et al. 2008), and that an increase in growth rate resulted in earlier puberty (Abeni *et al*., 2019). However, authors do not agree on the influence of earlier calving on milk performance: some observed a negative influence, while others did not. Abeni *et al*. (2000) and Van Amburgh *et al*. (1998) concluded that calving earlier than 23 mo is associated with lower milk yields and lower milk fat content; however, it also results in a higher milk protein content. They also concluded that earlier calving results in a decrease in reproductive performance. In a more recent study, Krpálková et al. (2014) observed that age at first calving had no influence on milk yields of primiparous cows, except for those during the first 100 d of lactation. They also observed the highest milk yield for the second and third lactation of heifers that first calved at 23 mo of age. In the present study, a negative influence was observed only at the start of the first lactation, but not for all of it. No data were available for later lactations. Van De Stroet *et al*. (2016) observed that primiparous cows that had consumed more starter feed as calves tended to have higher peak milk yields during first lactation than those that had consumed less. However, higher calf growth rates were not significantly related to future milk yield, but were related to higher BW of lactating cows and higher odds of surviving to first lactation. When lactation was corrected for BW, no difference in milk yield or composition was observed, regardless of the feeding strategy during the rearing period.

Decreasing the age of first calving is an effective way to decrease the length of the non-productive period during rearing. First calving at *ca*. 24 mo appears optimal for profitable production (Mourits *et al*., 1999b; Ettema and Santos; 2004; Shamay *et al*., 2005). In a meta-analysis of results of 100 herds, Mohd Nor *et al*. (2013) estimated that heifers that first calved at 24 mo produced a mean of 7 164 kg of milk per 305 d, and calving one mo earlier resulted in 143 kg less milk per 305 d. In the present study, younger heifers produced less milk during the first part of lactation, but the total milk yield per 305 d did not differ. The decrease in milk yield was similar (134 kg less per 305 d), albeit not significantly different, when age at first calving decreased from 24.8 to 23.8 mo of age. Age at first service had no effect on fertility. In a previous study on puberty attainment in the 2011-12 cohort, we observed that most heifers reached puberty before oestrus synchronisation, at aa mean age of 10.3 ± 2.2 mo (6.2-14.4 mo) and a mean BW of 296 ± 40 kg (224-369 kg; Abeni *et al*., 2019). ID2 heifers reached puberty one month earlier than SD and ID1 heifers. The onset of puberty at 9-10 mo or less meant that 3 or 4 oestrous cycles occurred before insemination, which is generally consistent with acceptable fertility results in many domestic species (Lin *et al*., 1986; Byerley *et al*., 1987; Robinson, 1990; Le Cozler *et al*., 1999). Regardless of calving strategy, decreasing the age of puberty and, consequently, the age of first service, is an effective way to shorten the non-productive period before calving. As Meyer *et al*. (2006) suggested, however, could reduce pre-pubertal mammary gland development by shortening the allometric phase of mammary gland growth and, in some cases, impair future milk production. Like its lack of effect on fertility in heifers, age at first calving did not influence fertility of primiparous cows during first lactation. Wathes *et al*. (2008) reported that fertility was optimised and maximum performance was maintained during first lactation when heifers first calved at 24-25 mo, although those that first calved at 22-23 mo had the best overall performance and longevity over 5 years, in partly because heifers with high fertility maintained high fertility as cows.

We also observed that at a similar feed allowance, early-calving heifers ate a similar amount of feed, produced less milk and ultimately were able to catch up in BW and development. As Krpalkova *et al*. (2014) reported, our results indicate that a feeding-rearing program that aims for first calving at less than 23 mo of age is a suitable option for successfully rearing Holstein heifers with optimal subsequent production and reproduction in a herd with suitable management. However, future studies are required to explore performances during the second and later lactations, as well as animal longevity.

## Ethics statement

Experimental work was conducted in accordance with French national legislation on the use of animals for research. Protocol agreement no. 00944-02 was received from French Ethical Committee n0.7.

## Data accessibility

Data are available online: https://doi.org/10.15454/CVKV1Z

## Supplementary material

Script and codes are available online: https://doi.org/10.15454/CVKV1Z

## Acknowledgements

The authors thank the technical staff of the INRA experimental farm of Méjusseaume for their commitment in taking care of the animals and making sure the experiment ran smoothly.

This preprint has been reviewed and recommended by Peer Community In Animal Science (https://doi.org/10.24072/pci.animsci.100002)

## Appendix

Appendix Fig.1 can be found together with the preprint on biorxiv: https://doi.org/10.1101/760082

## References

Abeni F, Calamari L, Stefanini L, Pirlo G 2000. Effects of daily gain in pre-and postpubertal replacement dairy heifers on body condition score, body size, metabolic profile, and future milk production. Journal of Dairy Science 83, 1468–1478.

Abeni F, Petrera F, Le Cozler Y 2019. Effects of feeding treatment on growth rates, metabolic profiles, and age at puberty, and their relationships in dairy heifers. Animal, 13(5):1020–1029.

Agabriel J, Meschy F 2007. Alimentation des veaux et génisses d’élevage. In Alimentation des bovins, ovins et caprins. Editions Quae, Versailles, chapitre 4, pp 75–87.

Bach A, Ahedo J 2008. Record keeping and economics of dairy heifers. Veterinary Clinics of North America Food Animal Practice, 24, 117–138.

Bach A 2011. Associations between several aspects of heifer development and dairy cow survivability to second lactation. Journal of Dairy Science, 94, 1052–1057.

Bartlett KS, McKeith FK, Van de Haar MJ, Dahl GE, Drackley JK 2006. Growth and body composition of dairy calves fed milk replacers containing different amounts of protein at two feeding rates. Journal of Animal Science, 84, 1454–1467.

Bazin S, Augeard P, Carteau M, Champion H, Chilliard Y, Cuylle G, Disenhaus C, Durand G, Espinasse R, Gascoin A, Godineau M, Jouanne D, Ollivier O, Remond B 1984. Grille de notation de l’état d’engraissement des vaches pie-noires. Institut Technique de l’Elevage Bovin, Paris, France.

Boulton AC, Rushton J, Wathes DC 2017. An empirical analysis of the cost of rearing dairy heifers from birth to first calving and the time taken to repay these costs. Animal, 11, 1372–1380.

Brickell JS, McGowan MM, Wathes DC 2009. Effect of management factors and blood metabolites during the rearing period on growth of dairy heifers on UK farms. Domestic Animal Endocrinology, 36, 67–81.

Byerley DJ, Staigmiller RB, Berardinelli JG, Short RE 1987. Pregnancy rates of beef heifers bred either on pubertal or third oestrus. Journal of Animal Science, 65, 645–650.

Cutullic, E, Delaby L, Gallard Y, Disenhaus C 2011. Dairy cows’ reproductive response to feeding level differs according to the reproductive stage and the breed, Animal, 5, 731–740.

Esslemont R, Kossaibati M 1997. The cost of respiratory diseases in dairy heifer calves. The Bovine Practitioner 33, 174–178.

Ettema JF, Santos EP 2004. Impact of age at calving on lactation, reproduction, health, and income in first-parity Holsteins on commercial farms. Journal of Dairy Science 87, 2730–2742.

Hammon HM, Schiessler G, Nussbaum A, Blum JW 2002. Feed Intake Patterns, Growth Performance, and Metabolic and Endocrine Traits in Calves Fed Unlimited Amounts of Colostrum and Milk by Automate, Starting in the Neonatal Period. Journal of Dairy Science, 85, 3352–3362

Hill T M, Bateman HG, Aldrich JM, Schlotterbeck RL 2010. Effect of milk replacer program on digestion of nutrients in dairy calves. Journal of Dairy Science, 93, 1105–1115.

INRA 2007. Alimentation des bovins, ovins et caprins – besoins des animaux – Valeurs des aliments – Tables INRA 2007. Edition Quae, Versailles, France, 307 pages.

INRA, 2018. Alimentation des ruminants. Editions Quae, Versailles, France, 728 p.

Jasper J, Weary DM 2002. Effects of ad libitum milk intake on dairy calves. Journal of Dairy Science, 85, 3054–3058.

Johnson KF, Vinod Nair R, Wathes DC 2019. Comparison of the effects of high and low milk-replacer feeding regimens on health and growth of crossbred dairy heifers. Animal Production Science, 59, 1648–1659.

Khan MA, Weary DM, von Keyserlingk MAG 2011. *Invited review:* Effects of milk ration on solid feed intake, weaning, and performance in dairy heifers. Journal of Dairy Science, 94, 1071–1081.

Krpálková L, Cabrera VE, Kvapilík J, Burdych J, Crump P 2014. Associations between age at first calving, rearing average daily weight gain, herd milk yield and dairy herd production, reproduction, and profitability. Journal of Dairy Science, 97, 6573–6582.

Le Cozler Y, Ringmar-Cederberg E, Johansen S, Dourmad JY, Neil M, Stern S, 1999. Effect of feeding level during rearing and mating strategy on performance of Swedish Yorkshire sows. 1. Growth, puberty and conception rate. Animal Science, 68, 355–363.

Le Cozler Y, Lollivier V, Lacasse P, Disenhaus C 2008. Rearing strategy and optimizing first-calving targets in dairy heifers: a review. Animal, 2, 1393–1404.

Lin CY, McAllister AJ, Batra TR, Lee AJ, Roy GL, Vesely JA, Wauthy JM, Winter KA 1986. Production and reproduction of early and late bred dairy heifers. Journal of Dairy Science, 69, 760–768.

Meyer MJ, Capuco AV, Ross DA, Lintault LM, Van Amburgh ME 2006. Development and nutritional regulation of the prepubertal heifer mammary gland: I. Parenchyma and fat pad mass and composition. Journal of Dairy Science 89, 4289–4297.

Mohd Nor N, Steeneveld W, van Werven T, Mourits MCM, Hogeveen H 2013. First-calving age and first-lactation milk production on Dutch dairy farms. Journal of Dairy Science, 96, 981–992.

Morrison SJ, Wicks HCF, Fallon RJ, Twigge J, Dawson LER, Wylie ARG, Carson AF 2009. Effects of feeding level and protein content of milk replacer on the performance of dairy herd replacements. Animal, 3, 1570–1579.

Mourits MCM, Huirne RBM, Dijkhuizen AA, Kristensen AR, Galligan DT 1999. Economic optimization of dairy heifer management decisions. Agricultural Systems, 61, 17–31.

Petersson KJ, Gustafsson H, Strandberg E, Berglund B 2006. Atypical progesterone profiles and fertility in Swedish dairy cows. Journal of Dairy Science, 89, 2529–2538.

Pirlo G, Capelleti M, Marchetto G 1997. Effects of energy and protein allowances in the diets of prepubertal heifers on growth and milk production. Journal of Dairy Science, 80, 730–739.

Quigley JD, Wolfe TA, Elsasser TH 2006. Effects of additional milk replacer feeding on calf health, growth, and selected blood metabolites in calves. Journal of Dairy Science, 89, 207–216.

Robinson JJ 1990. Nutrition in the reproduction of farm animals. Nutrition Research Reviews, 3, 253–276.

R Core Team 2019. R: A Language and Environment for Statistical Computing. R Development Core Team, Vienna, Austria.

Shamay A, Homans R, Fuerman Y, Levin I, Barash H, Silanikove N, Mabjeesh SJ 2005. Expression of albumin in nonhepatic tissues and its synthesis by the bovine mammary gland. Journal of Dairy Science, 88, 569–576.

Smith JM, Van Amburgh ME, Diaz MC, Lucy MC, Bauman DE 2002. Effectof nutrient intake on the development of the somatotropic axis and itsresponsiveness to GH in Holstein bull calves. Journal of Animal Science, 80, 1528–1537.

Soberon F, Raffrenato E, Everett RW, van Amburgh ME 2012. Preweaning milk replacer intake and effects on long-term productivity of dairy calves. Journal of Dairy Science, 95, 783–793.

Tozer PR 2000. Least-cost ration formulations for Holstein dairy heifers by using linear and stochastic programming. Journal of Dairy Science 83, 443–451.

Van Amburgh ME, Galton DM, Fox DG, Bauman DE, Chase LE, Erb HN, Everett RW 1998. Effects of three prepubertal body growth rates on performance of Holstein heifers during first lactation. Journal of Dairy Science, 81, 527–538.

Van Amburgh ME, Drackley J 2005. Current perspectives on the energy and protein requirements of the preweaned calf. Chapter 5 in Calf and Heifer Rearing. P.C. Garnsworthy, ed. Nottingham University Press, Nottingham, UK.

Van De Stroet DL, Calderón Díaz JA, Stalder KJ, Heinrichs AJ, Dechow CD, 2016. Association of calf growth traits with production characteristics in dairy cattle. Journal of Dairy Science 99, 8347–8355.

Wathes DC, Brickell JS, Bourne NE, Swali A, Cheng Z 2008. Factors influencing heifer survival and fertility on commercial dairy farms. Animal, 2, 1135–1143.

Wathes DC, Pollott GE, Johnson KF, Richardson H, Cooke JS 2014. Heifer fertility and carry over consequences for lifetime production in dairy and beef cattle. Animal 8 (suppl. 1), 91–104.

