## Appendix Fig.1 for "Effects of feeding treatment on growth rate and performance of primiparous Holstein dairy heifers"

Supplementary files

Figure 1: monitor morphological traits during rearing and first lactation, according to age at first calving

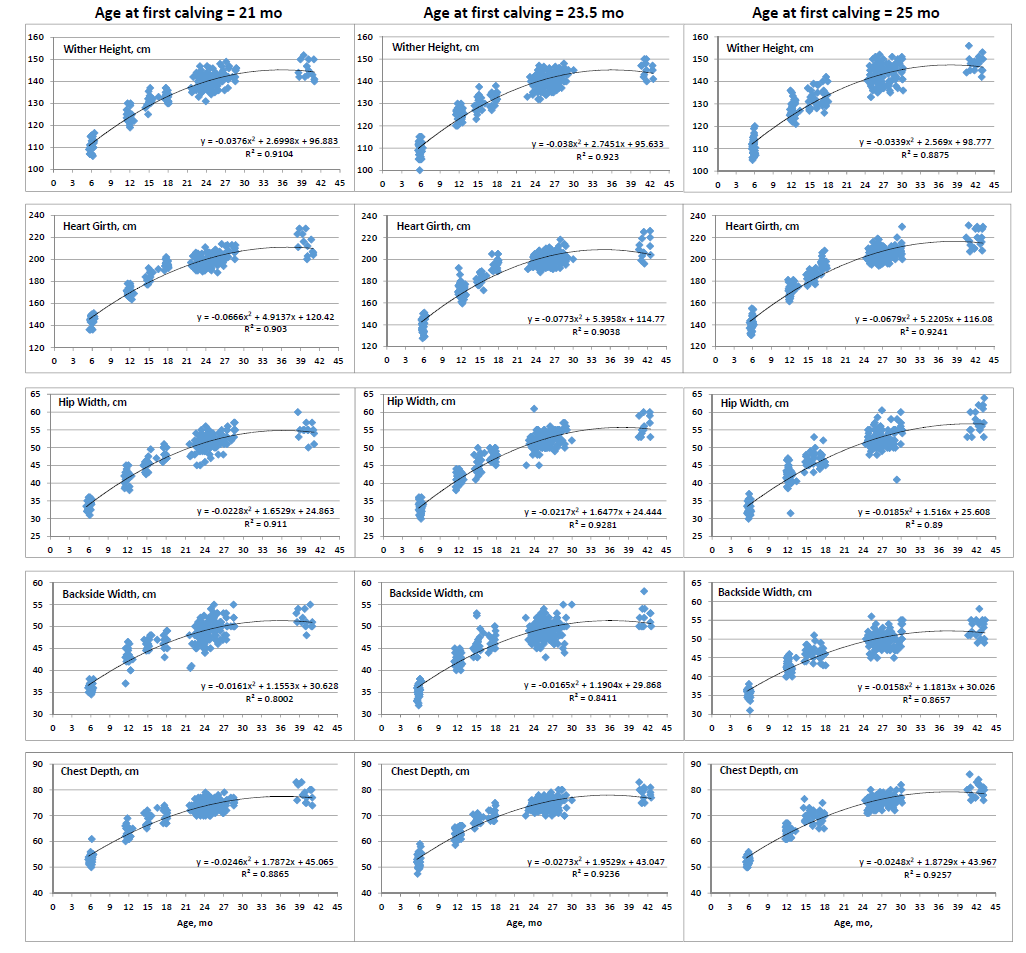
